# Ramifications of the diverse mRNA patterns in *Acanthamoeba royreba*

**DOI:** 10.1101/2020.05.17.100792

**Authors:** Richard Tyndall, Ibne Ali, Anthony Newsome

## Abstract

Free-living amoebae are distributed worldwide and can be found in a variety of environments. While most Acanthamoebae have been isolated from soil and water, *A. royreba* and *A. culbertsoni* were isolated from mammalian cell cultures. *A. royreba’s* isolation from placental and tumoral tissue, its unusual ability to grow in mammalian cell culture media at 35°C, and the presence of a primitive centriole, prompted us to investigate the potential for mammalian information in *A. royreba.* While some soil amoeba contain small amounts of fungal and bacterial information, presumably from the microbes they phagocytosed, the informational content of *A. royreba,* in some instances, was very different with > 70% of the mRNA being non-amoebic. Here we show that the proteins and mRNA content associated with *A. royreba* are extremely diverse and represent multiple Kingdoms, Orders, Phyla, and Genera from around the globe. The information in *A. royreba,* such as placental proteins from numerous mammals, the preponderance of non-amoebic mRNA, and its ability to tolerate harsh environments including Megarad irradiation, leads to a discussion regarding the possible role of this amoebae as an immunological gatekeeper protecting fetal or malignant tissue from destruction and its potential as a vehicle for panspermia.

## Introduction

*Acanthamoeba* was first described by Castellani [1] and represents single-cell eukaryotes existing as either cellular trophozoites (25-40 μm) or, under adverse conditions (dessication, lack of food, and extreme pH or temperature fluctuations) [2, 3], as dormant cysts (13-20 μm). The cysts are known to be resistant to antibiotics, the effects of chlorine, and very low temperatures [4–6], and have been shown to maintain viability for over 20 years [7]. *Acanthamoeba* spp. have been placed into three groups (I, II, and III) based on the morphology and size of their cysts [8, 9]. The 21 known *Acanthamoeba* spp. are ubiquitous, have been isolated from diverse environments including soil, water, compost, marine sediments, shellfish beds, and sludges [10–16]. They have also been isolated from fish, reptiles, and amphibians [17, 18]. Two group III species, specifically *A. culbertsoni* and *A. royreba,* have been isolated from mammalian cell cultures [19, 20]. Several have been implicated in a variety of diseases such as keratitis [21], encephalitis [22], and sinusitis [23].

Relative to a study we were undertaking regarding gallium-67 (^67^Ga) uptake in mammalian placentas and tumors [24, 25], we were interested in pursuing the interaction of ^67^Ga with cultured human choriocarcinoma cells. Consequently, we obtained a culture of BeWo cells from Dr. Pattillo at the Medical College of Wisconsin (Milwaukee, Wisconsin, USA). This was the first human trophoblastic endocrine cell type to be maintained in continuous culture. We grew the culture in a mammalian cell culture medium (Plus I) with 10% added fetal calf sera (FCS). Shortly after obtaining this culture, events occurred which necessitated our laboratory being vacated for several weeks. In the interim, the BeWo cells had not received any media changes and subsequently experienced detrimental effects. While a majority of the BeWo cells were dead and floating upon our return, some were still attached and appeared viable. Surprisingly, some amoeboid motile cells were also observed intermixed with the few surviving BeWo cells. We subcultured the living amoeboid cells and sent them to Dr. Willaert at the Department of Protozoology, Institute of Tropical Medicine, Antwerp, Belgium. He observed that the locomotion of the cells was by protrusion of the cell cytoplasm involving the formation of pseudopodia with the creation and resorption of numerous spiked filiform projections. Under microscopic examination he determined that the amoebae were approximately 12–25 μm in length with a diameter of 10–15 μm. The amoebae also had a primitive centriole, relatively small nucleus with a single centrally located nucleolus, and abundant vacuoles. When the cells were placed in water they did not flagellate and when placed in saline or in media without added sera, the amoebae formed cysts typical of Acanthamoebae. Ultimately he determined that the amoeboid cells were a new species: *Acanthamoeba royreba* [20, 26].

While uncommon, ‘infection’ or ‘contamination’ of cultured mammalian cells (primary monkey kidneys cells, HeLa cells, and chick embryo cells) with free-living *Acanthamoeba* has been reported in the literature [19, 27–29], with the first of these reports discussing the infection of cultured monkey kidney cells by an unidentified amoebae [30].

In light of the occasional unplanned sporadic isolation of free-living amoeba from cultured mammalian cells, including the unexpected isolation of *A. royreba* from BeWo cells, we attempted to deliberately induce the emergence of Acanthamoebae from various mammalian cell cultures by starvation. The untransformed cell lines tested included human W1-38 and mouse A-31 cell cultures. Malignant cell lines tested included rat hepatoma H35 and virus-transformed A31 Balb-C (KBALB) mouse cultures.

Plus I supplemented with 10% FCS was used as the base medium for the cell cultures which were starved (no medium exchanges) for 4–20 weeks. Visual and electron microscopic examination of the cultures after the starvation period did not reveal any evidence of amoebic cysts or trophozoites. However, while the addition of American Type Culture Collection (ATCC, Manassas, VA, USA) casitone medium #712 (typically used for *Acanthamoeba* cultures) to the flasks containing the starved cultures usually produced no growth. However, a few of the KBALB and H35 flasks did show the appearance of amoeboid cells after several attempts. The flasks which contained only the Plus I medium and FCS did not reveal the appearance of amoeboid cells nor did any of the A-31 or W1-38 cell cultures. Similar to the *A. royreba* culture isolated from the BeWo culture, the amoeboid cells from the KBALB and H35 cultures had characteristics (bulls-eye nucleolus, pseudopods, formed cysts, did not flagellate, and had vacuoles) typical of *Acanthamoeba* and were confirmed as such using immunological and serological assays. Dr. Willaert subsequently identified these isolates as *A. culbertsoni* [26].

All of the isolations of *Acanthamoeba* were difficult and required careful manipulation, observation, and the use of selective culture media. Ultimately, we felt that a more sophisticated approach to detect *Acanthamoeba* in placental or malignant tissue would be by the use of monoclonal antibodies. With some difficulty, a monoclonal antibody against *A. royreba* was produced [31].

## Results and discussion

*A. culbertsoni* was first isolated from monkey kidney cells [19], and the isolation of *A. royreba* occurred when human choriocarcinoma cells [32], were stressed by not changing the liquid culture media for several weeks. Similarly, when we deliberately starved separate cultures of malignant rat and mouse cells, we were also able to isolate cultures of *A. culbertsoni* [19]. The identification of *A. royreba* and *A. culbertson*i from four separate axenic mammalian cultures prompted further investigations to understand the possible interaction between *Acanthamoeba* and other prokaryotes/eukaryotes. During our initial foray into these studies, we were curious whether phagocytic *A. royreba* would support replication of *Legionella* bacteria in manner similar to human macrophages; therefore, we deliberately infected *A. royreba* with an axenic culture of *Legionella pneumophila* [33], a Gram-negative bacterium that causes legionellosis [34]. While copious amounts of *Legionella* were replicated in *A. royreba* and subsequently cultured on Buffered Charcoal Yeast Extract medium, small amounts of diverse, somewhat unusual bacterial also grew on the *Legionella*-specific culture medium and tended to have unusual properties (highly pigmented and the ability to produce biogenic crystals) which differed from their conventional counterparts [35]. These bacteria were not culturable from the amoebae in the absence of the stress caused by the *Legionella* infection. This led to subsequent experiments using a ‘sterile stressor’, 1 Megarad gamma radiation, to further stress the amoeba with even more provocative results. As with the *Legionella* infection, previously unculturable bacteria could now be grown post-radiation [35]. In addition to biogenic crystal and pigment formation, many also produced extremely powerful biodispersants (capable of emulsifying mercury) and showed a diminished ability to ferment common sugars. Perhaps most interesting was that not only did the amoebic trophozoites survive the radiation dosage, but the bacteria that arose after gamma radiation were also radiation resistant! While stressing *A. royreba* resulting in the appearance of culturable bacteria mimics the stressing of the choriocarcinoma cells to yield culturable *A. royreba* in a Lego-like-linkage, the more interesting aspect of the radiation experiment was the ‘lethal/sterilizing’ Megarad dosage of gamma radiation used.

Having studied the interaction of *A. royreba* with *Legionella* which revealed the presence of additional prokaryotic bacteria, we then explored the possible sequestration of eukaryotic information in this amoebae since it exhibited some enigmatic mammalian characteristics. Not only was it isolated from human choriocarcinoma cells, but it could grow in several media used specifically for mammalian cell cultures and at temperatures as high as 35° C. Additionally, the cysts from this species could survive temperatures as low as −85^0^ C. Unlike most other *Acanthamoeba*, *A. royreba* and *A. culbertsoni* have primitive centrioles [20], an organelle which plays a major role in microtubule organization during cell division in other eukaryotes. Most importantly, monoclonal antibody production against *A. royreba*, while easily accomplished in male mice, could not be accomplished using female mice. Interestingly, monoclonal antibodies that were produced against *A. royreba* interacted with a subpopulation of cells obtained from a monkey placental tissue (unpublished observations).

Initially, *A. royreba* was analyzed to detect the presence of mammalian proteins. Shotgun protein analysis was performed by Bioproximity (Manassas, VA, USA) [39–41], and the results indicated hundreds of mammalian and human-specific proteins ranging from lymphocytes, erythrocytes, tumor rejection antigens, and sperm-binding proteins as well as several pregnancy-specific glycoproteins. While no one type commanded a high percentage of the total protein content, the diversity of these proteins was intriguing (Table 1). These results were interesting in that the protein information expressed in *A. royreba* contrasted with the genetic information in *A. castellani* as eloquently shown by Clarke et al. [36]. The genome of *A. castellani* showed the presence of predominantly Acanthamoeba genes and bacterial and fungal information which amounted to approximately 2.9% of the genome, presumably from ingestion of these microbes, and could represent lateral gene transfer (LGT). LGT, or horizontal gene transfer [37, 38], is the transmission of genes between individual cells (from species to species) and can generate new gene sequences through transformation, transduction, and conjugation.

**Table 1.** Representative protein groups isolated from *Acanthamoeba royreba.*

| **Archaea (0%)** | **Bacteria (37.69%)** | **Eukarya (62.31%)** |
| --- | --- | --- |
|  | 15 Phyla<sup>1</sup> | Fungi – 3 Phyla<sup>2</sup> |
|  |  | Animalia – 4 Orders<sup>3</sup> |
|  |  | Protista<sup>4</sup> |

<sup>1</sup> representative habitats include: marine, freshwater, brackish, sludge, sand, gut flora, soil
<sup>2</sup> includes lichens
<sup>3</sup> representative mammals include: humans, placental mammals, shrews, bats, even-toed hoofed animals, new world monkeys, African ruminant animals, cattle
<sup>4</sup> includes 4 spp. of *Acanthamoeba* (*A. castellanii*: 99.99%) and Proteobacteria

Subsequently, *A. royreba,* axenically cultured in a proteose peptone-yeast extract-glucose (PYG) medium with 40 μg gentamicin/ml [42], was tested to determine its messenger RNA (mRNA) content by both Quick Biology (Pasadena, CA, USA) and My Genomics (Atlanta, GA, USA).

The diversity shown in the protein analysis of *A. royreba* was further confirmed and greatly magnified in the content of the mRNAs (Tables 2 and 3, and Figures 1 and 2). Duplicate frozen cultures of *A. royreba* were thawed and grown at 37° C. The results showed that after 2 weeks approximately 25% of the mRNA profile was non-amoebic (Figure 1) which increased to approximately 70% after 2 months (Figure 2), indicating a possible time- and/or temperature-dependent genetic expression component. While not the norm, growing *A. royreba* at 37° C may be the road best traveled for revealing pivotal mRNA profiles. While less than half of the mRNA in the cells cultured for 2 months represented characteristics specific to *Acanthamoeba*, the remaining spectrum of mRNA content was very diverse and represented fungal, chordata, arthropoda, and bacterial communities. As noted in the protein analysis, approximately 3% of the mRNA was mammalian (Tables 2 and 3, and Figure 2).

**Figure 1.**
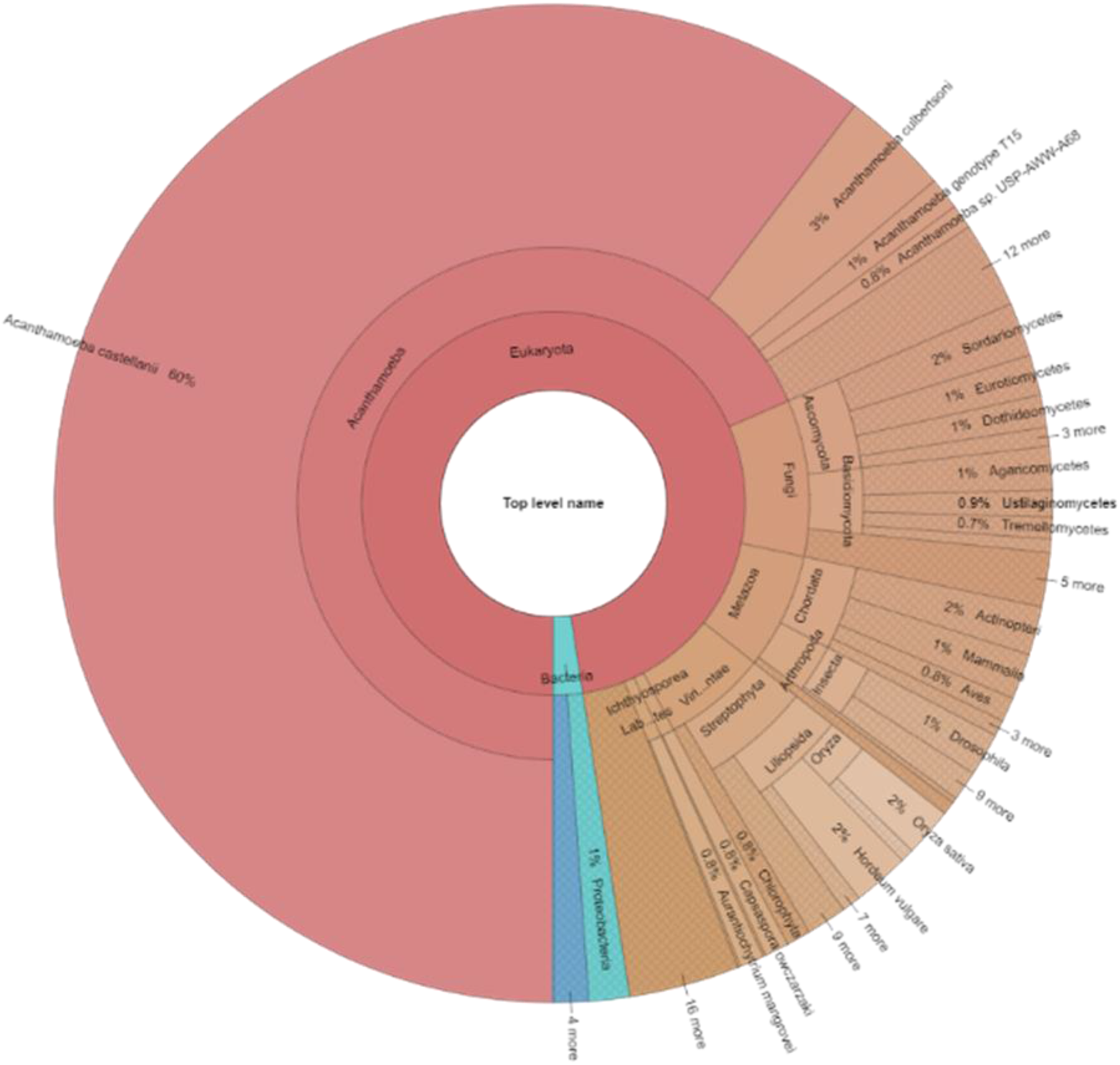
mRNA analysis of A. royreba grown for 2 weeks prior to analysis showing diversity of taxa.

**Figure 2.**
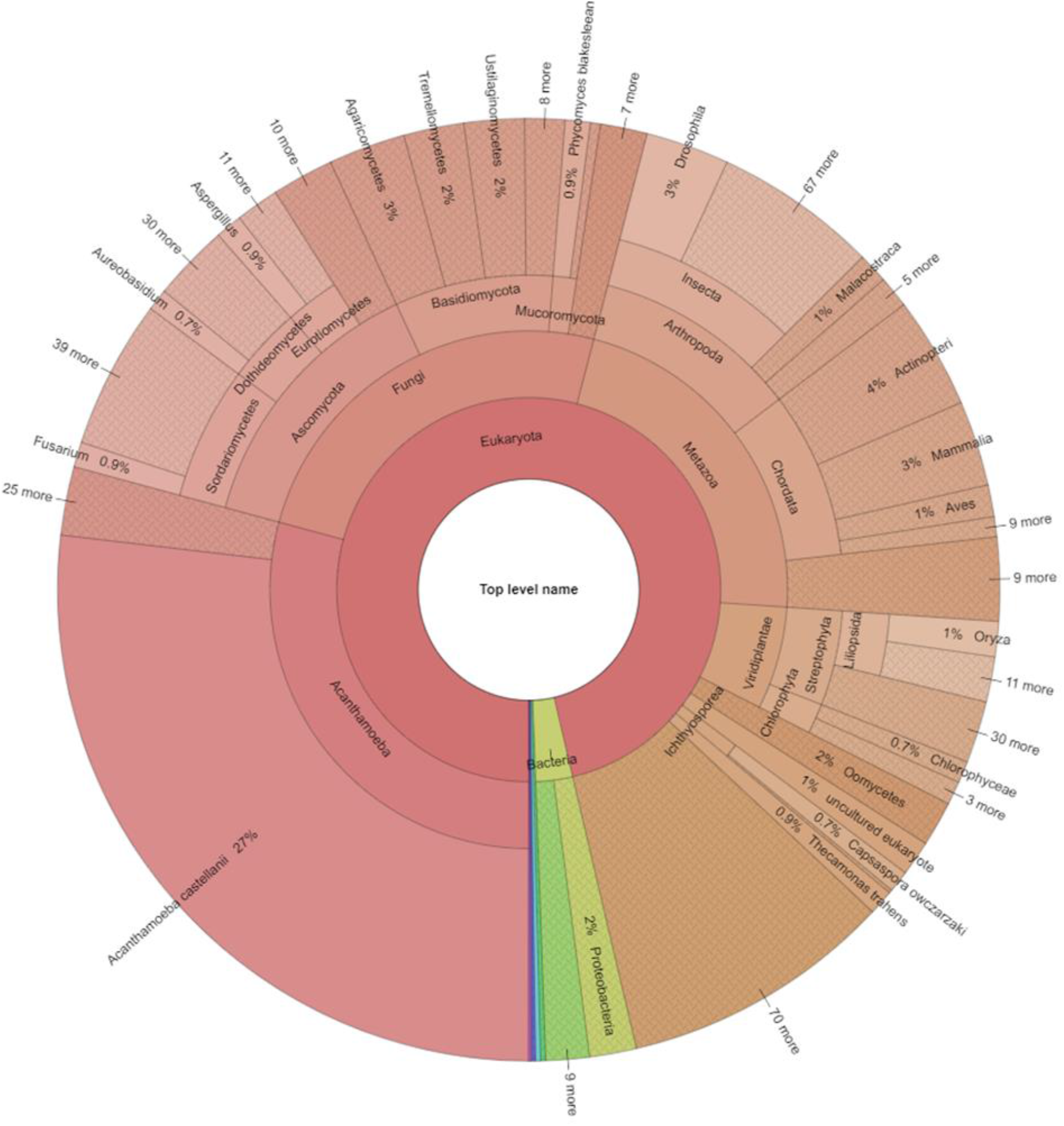
mRNA analysis of A. royreba grown for 2 months prior to analysis showing extreme diversity of taxa.

**Table 2.**
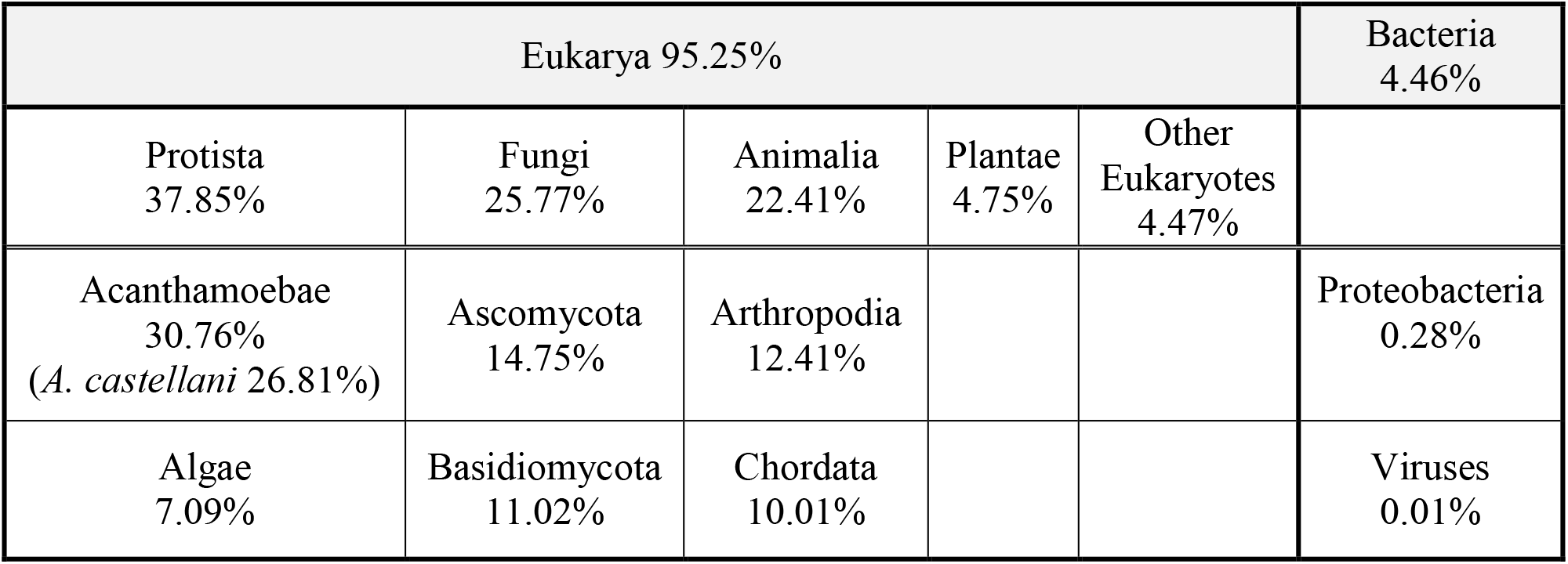
mRNA sequences from various taxa identified in *Acanthamoeba royreba.*

| Eukarya 95.25% |  |  |  |  | Bacteria<br>4.46% |
| --- | --- | --- | --- | --- | --- |
| Protista<br>37.85% | Fungi<br>25.77% | Animalia<br>22.41% | Plantae<br>4.75% | Other<br>Eukaryotes<br>4.47% |  |
| Acanthamoebae<br>30.76%<br>(*A. castellani* 26.81%) | Ascomycota<br>14.75% | Arthropodia<br>12.41% |  |  | Proteobacteria<br>0.28% |
| Algae<br>7.09% | Basidiomycota<br>11.02% | Chordata<br>10.01% |  |  | Viruses<br>0.01% |

**Table 3.** Examples of the extreme diversity^1^ of mRNA identified in *Acanthamoeba royreba.*

|  |  |  |
| --- | --- | --- |
| Algae | 99 different Genera | Red, green, flagellated |
| Fungi | 360 different Genera | Mushrooms, yeasts, molds |
| Bacteria | 67 different Genera | Pathogens, nonpathogens, soil, water, hot springs |
| Insects | 120 different Genera | Mosquitos, flies, moths, butterflies, ants, wasps, bees, beetles, planthopper, bed bugs, mites, aphids, ticks |
| Fish | 41 different Genera | Coelacanth, whitefish, sablefish, minnow, lancelets, herrings, pupfish, carp, danios, pike, cichlid, catfish, croakers, gars, icefish, mollies, damselfish |
| Marine organisms and crustaceans | 26 different Genera | Crabs, lobsters, squid, sperm whale, sea squirt, prawn, sea slug, oysters, barnacles, tube worms, sponges |
| Mammals and amphibians | 52 different Genera | Mouse, camel, elephant, goat, alligator, lizard, Seriema, guinea pig, boar, turtles, crocodiles, bats, gecko, gorilla, squirrel, Weddell seal, turkey, hamster, lemurs, ferret, mole-rat, frogs, degu, aardvark, rabbit, baboons, shrews |
| Crops | 22 different Genera | Rice, corn, cucumber, soybean, apples, pears, radishes, castor bean, sesame |
| Birds | 11 different Genera | Pigeons, falcon, woodpeckers, starlings |
| Trees | 21 different Genera | Gum tree, locust tree, walnut tree, date palm, poplar, tropical evergreens |
| Plants | 17 different Genera | Rose, lupine, various grasses |

<sup>1</sup>Nearly all Genera included more than 1 species

Since many Acanthamoebae live primarily in soil and water, ingest a variety of fungal and bacterial microbes, and contain bacterial endosymbionts, it is understandable that the amoebae contain mRNAs indicative of these organisms. However, the fact that there is a species, *A. royreba*, which contains mRNA specific to a wide and diverse spectra within multiple Kingdoms, Phyla, and Genera is both puzzling and fascinating and may have profound evolutionary implications.

The questions these experiments raise have profound implications in evolutionary biology, physiology, environmental science, medicine, exobiology, and paleobiology, to mention just a few. Several admittedly speculative implications arise from *A. royreba* being an immense reservoir of global genetic information:

1. Is there some analogy to be drawn from the relationship between mammalian cells and an amoebae and the bacterial origin of mitochondria [43]?
2. *A. royreba* was originally isolated from a malignant human placental trophoblast (i.e., choriocarcinoma), contains placental proteins from various mammals, and was not recognized as foreign by female mice, but was recognized as such by male mice. Also, the antibodies made against *A. royreba* did react with some cells from a Rhesus monkey placenta. Therefore, could these observations indicate that this amoebae exists as an endosymbiont in placental or tumoral tissue? Could it possibly play a crucial role in the womb becoming an immunologic sanctuary that permits the presence of a foreign fetus without rejection? Can this unusual property be extrapolated to explain the tolerance of the immune system to fetal or tumoral tissue? Interestingly, when *A. royreba* was placed in a co-culture of human red and white blood cells, the amoebae preferentially destroyed the white cells [19]. Similarly, the study by Chi et al. [28], also showed that the *Acanthamoeba* they isolated from monkey kidney cells would specifically attack and enucleate chick erythrocytes, but would ignore guinea pig erythrocytes. These observations indicate a mechanism whereby amoebae can ‘identify’ different cell types. Obviously a placenta-based amoebic ‘immunologic gate keeper’ would want to allow entry to red blood cells, but not to potentially harmful white blood cells.
3. *A. royreba* can tolerate temperatures of −85° C, MegaRad doses of gamma radiation, form cysts that contain and safeguard a vast spectrum of genetic information, and harbor or spawn numerous bacteria or bacteria-like organisms with unusual properties in an apparent symbiotic-like relationship allowing both to survive harsh environments. Can it therefore be a candidate as a vehicle for panspermia thereby shedding light on the transfer of life [44, 45]? Is it also imaginable that an entity such as *A. royreba* could, perhaps encased in asteroids, comets, or space dust, traverse many light years and seed planets in the cosmos – including Earth?

Obviously, more intense light needs to be applied to this initial study. Now we look through the glass darkly, but the images are still very exciting.

## Materials and methods

Protein and mRNA analyses:

Shotgun protein analysis was performed by Bioproximity: 9385 Discovery Blvd

Manassas, VA 20109;

mRNA analyses was performed by Quick Biology: 2265 E Foothill Blvd., Pasadena, CA, 91107 and My Genomics; Atlanta, GA, USA;

RNA integrity was checked using an Agilent Bioanalyzer 2100 (Agilent, Santa Clara, CA, USA); only samples with clean rRNA peaks were used for further experiments. The RNA-Seq Library was prepared according to KAPA Stranded mRNA-Seq poly(A) selected kits with 201-300 bp insert sizes (KAPA Biosystems, Wilmington, MA, USA) using 250 ng total RNA as an input. Final library quality and quantity were analyzed using an Agilent Bioanalyzer 2100 and a Life Technologies Qubit 3.0 Fluorometer (ThermoFisher Scientific, Waltham, MA, USA). 150 bp paired-end reads were sequenced on an Illumina HiSeq 4000 (Illumnia Inc., San Diego, CA). 10,000 random reads were used for the blast searching against the NCBI nr database. A custom perl script was used for the taxonomy analysis based on the blast result. Krona was used to generate multi-layered pie charts.

## Methods regarding the growth of discussed organisms

Amoebae were grown following the procedure of Willaert et al. [20].

Choriocarcinoma cells were cultured as described in Pattile et al. [32].

Monoclonal antibody production in mice used the procedure of Kennel et al. [31].

Gamma irradiation procedures and the growth and isolation of endosymbionts followed the protocol outlined in Vass et al. [35].

*Legionella* were cultured as described in Tyndall et al. [33] and Lau et al. [34].

*A. royreba*, was axenically cultured in a proteose peptone-yeast extract-glucose (PYG) medium with 40 μg gentamicin/ml [42].

## Acknowledgments

We wish to thank Audrey Nicholson who first detected A. royreba in the choriocarcinoma cell culture and Shantanu Roy for assistance in sample preparation. Quick Biology, Pasadena, CA, provided Figures 1 and 2.

## Disclaimer

The findings and conclusions in this report are those of the authors and do not necessarily represent the official position of the Centers for Disease Control and Prevention.

